# Amyloidogenic Propensity of Self-Assembling Peptides and their Adjuvant Potential for use as DNA Vaccines

**DOI:** 10.1101/2022.09.09.507367

**Authors:** Paresh C. Shrimali, Sheng Chen, Rachel Dreher, Matthew K. Howard, Jeremy Buck, Darren Kim, Jai S. Rudra, Meredith E. Jackrel

## Abstract

*De novo* designed peptides that self-assemble into cross-β rich fibrillar biomaterials have been pursued as an innovative platform for the development of adjuvant- and inflammation-free vaccines. However, they share structural properties similar to amyloid species implicated in neurodegenerative diseases, which has been a long-standing concern for their translation. Here, we comprehensively characterize the amyloidogenic character of the amphipathic self-assembling cross-β peptide KFE_8_, compared to pathological amyloid and amyloid-like proteins α-synuclein (α-syn) and TDP-43. Further, we developed plasmid-based DNA vaccines with the KFE_8_ backbone serving as a fibrillizing scaffold for delivery of a GFP model antigen. We find that expression of tandem repeats of KFE_8_ is non-toxic and can be efficiently cleared by autophagy. We also demonstrate that synthetic KFE_8_ nanofibers do not cross-seed amyloid formation of α-syn in mammalian cells compared to α-syn preformed fibrils. In mice, vaccination with plasmids encoding the KFE_32_-GFP fusion protein elicited robust immune responses, inducing production of significantly higher levels of anti-GFP antibodies compared to soluble GFP or α-syn tagged GFP. Antigen-specific CD8^+^T cells were also detected in the spleens of vaccinated mice and cytokine profiles from antigen recall assays indicate a balanced Th1/Th2 response. These findings illustrate that cross-β-rich peptide nanofibers have distinct properties from those of pathological amyloidogenic proteins, and are an attractive platform for the development of DNA vaccines with self-adjuvanting properties and improved safety profiles.

## INTRODUCTION

Natural and *de novo* designed peptides that self-assemble into β-rich nanofibers have emerged as excellent building blocks for fabricating materials with a variety of biomedical applications[1]. Some of the simplest and most widely used synthetic peptides are amphipathic and comprised of alternating hydrophobic and polar/charged amino acids. Several such peptides have been shown to spontaneously associate to form fibrillar scaffolds in physiological buffers[1, 2]. Self-assembling peptide biomaterials have several key advantages. First, they only require the synthesis of a simple peptide building block which then self-associates to form higher order complex structures. Additionally, functional groups can be appended to the peptides’ *N*- or *C*-termini which are displayed in a multivalent fashion along the fibril surface, thereby imparting biofunctionality[3]. Numerous studies have demonstrated that self-assembling peptide biomaterials can support the growth of a variety of cell types, deliver therapeutic drugs, exert antimicrobial effects, serve as theranostics, and act as immune adjuvants in vaccines[1, 4, 5].

A range of self-assembling peptide sequences have been designed that adopt a variety of secondary structures[6]. A particularly promising design is based on the cross-β structure where β-strands form the fibril axis which is stabilized by non-covalent interactions such as hydrophobic interactions, π–π stacking, and electrostatic interactions[7]. While these cross-β rich scaffolds are highly promising as biomaterials, cross-β structures are also a defining characteristic of disease-relevant amyloid fibrils[8]. Indeed, accumulations of amyloid fibrils and pre-amyloid species are the pathological hallmark of many neurodegenerative disorders including Alzheimer’s disease and related dementias[9-11]. Despite their structural and morphological similarity to disease-associated amyloids, β-rich fibrillar biomaterials have been explored for multiple biological applications, without reports of their toxicity. We hypothesize that this is because the designed peptides aggregate extremely rapidly, and the pre-amyloid oligomeric species are likely the most toxic[12]. These differing kinetics likely bypass, or greatly decrease, the length of time that the nanofibers populate the highly toxic pre-amyloid oligomeric state. Another key feature of amyloids is their ability to seed further aggregation by nucleating amyloidogenesis[13]. Here, a small amount of amyloid fibrils can spontaneously enter cells and nucleate or ‘seed’ the amyloidogenesis of additional copies of monomer, thereby rapidly increasing amyloidogenesis in neighboring cells[13, 14]. It is crucial that any amyloid-inspired biomaterial be unable to seed the amyloid cascade of other amyloid-prone proteins. Therefore, it is crucial that the properties of cross β-sheet rich nanofibers be comprehensively characterized prior to their further application in these contexts[13, 15].

The KFE_8_ sequence (FKFEFKFE) is one of the best characterized cross-β sequences used for generating hydrogel biomaterials, with ample data pertaining to the effects of sequence length, pattern variation, and chiral substitutions on self-assembly and molecular packing[7, 16-19]. This sequence is amphipathic, enriched in Phe, and comprised of alternating apolar and charged residues. The two apolar faces associate to form a cross-β motif that forms the backbone of the fibrils. We have demonstrated that KFE_8_ is a powerful immune adjuvant for the development of vaccines and immunotherapies in multiple preclinical disease models[20-23]. To further apply KFE_8_-based vaccines, it is crucial that they first be comprehensively characterized. Here, to accelerate translational efforts, different length repeats of KFE_8_ (KFE_8_, KFE_16_, and KFE_32_) were expressed in a yeast model system and their toxicity and clearance mechanism was compared to that of the toxic amyloid-like protein TDP-43, which is implicated in amyotrophic lateral sclerosis (ALS) and frontotemporal dementia (FTD)[24]. We also investigated the seeding capacity of KFE_8_ nanofibers because it is known that some proteins can cross-seed, whereby fibrils of one protein can trigger fibrillization of a different amyloid-prone protein[14]. Here, we tested the capacity of KFE_8_ to cross-seed the aggregation of α-synuclein, a key protein implicated in Parkinson’s disease (PD), using a mammalian biosensor cell line[25, 26]. The successes we achieved in recapitulating the properties of the synthetic peptide nanofibers upon cellular expression suggest that this system might also be harnessed as a new form of DNA-based vaccine. Therefore, we generated lentivirus vectors expressing soluble GFP, α-syn-GFP or KFE_32_-GFP for expression in mammalian cells, with GFP serving as both an expression marker and a model antigen. Mice were vaccinated with plasmids encoding soluble GFP, α-syn-GFP or KFE_32_-GFP and antibody and cellular immune responses were evaluated. Our results demonstrate that despite their molecular, structural, and morphological similarity to pathological amyloids, self-assembling peptides have distinct properties, and are promising materials for biomedical applications including DNA-based vaccines.

## MATERIALS AND METHODS

### Yeast strains, Media, and Plasmids

All yeast were BY4741 or BY4741Δ*atg8*, acquired from the Yeast Knockout Collection[27]. pAG423GAL-ccdB-eGFP plasmid for yeast expression was obtained from Addgene[28]. Strain expressing TDP-43 has been described previously[29]. Yeast were grown in rich medium (YPD) or in synthetic media lacking the appropriate amino acids. Media was supplemented with 2% glucose, raffinose, or galactose. To construct KFE_8_ plasmids, repeating sequences of KFE_8_ interspersed with flexible glycine-serine linker regions were synthesized by IDT with flanking Gateway cloning sites. These fragments were then inserted into pAG423GAL-ccdB-eGFP[28] by Gateway cloning to generate KFE_8_ tagged with GFP at the C-terminus. Final sequences were: KFE_8_: MFKFEFKFEGSGSGSGS-(eGFP), KFE_16_: MFKFEFKFEGSGSGSGSFKFEFKFEGSGSGSGS-(eGFP), and KFE_32_: MFKFEFKFESGSGSGSGFKFEFKFEGSGSGSGSGFKFEFKFEGGSGSGSGFKFEFKFEGSG SGSGS-(eGFP).

### Yeast Transformation and Spotting Assays

Yeast were transformed according to standard protocols using polyethylene glycol and lithium acetate[30]. For the spotting assays, yeast were grown to saturation overnight in raffinose supplemented dropout media at 30ºC, normalized to A_600nm_ = 1.5, serially diluted, and spotted in duplicate onto synthetic dropout media containing glucose or galactose. Plates were analyzed after growth for 2-3 days at 30ºC. Each experiment was repeated three times.

### Immunoblotting and Microscopy

Yeast were grown and induced in galactose containing medium for 5h from overnight cultures supplemented with raffinose. Cultures were normalized to A_600nm_=0.6, 8 mL cells were harvested, treated in 0.1M NaOH for 5 min at room temperature, and cell pellets were then resuspended into 1x SDS sample buffer and boiled for 4min. Lysates were cleared by centrifugation at 14,000 rpm for 2min and then separated by SDS-PAGE (4-20% gradient, BioRad), and transferred to a PVDF membrane. Membranes were blocked in Odyssey Blocking Buffer (LI-COR). Primary antibody incubations were performed at 4ºC overnight. Antibodies used: anti-GFP monoclonal (Roche Applied Science), anti-TDP-43 polyclonal (Proteintech), and anti-PGK monoclonal (Invitrogen). Membranes were imaged using a LI-COR Odyssey FC Imaging system. For microscopy, strains were grown as for immunoblotting. After 15h induction at 30ºC, cultures were harvested and processed for microscopy. All imaging was performed using live cells treated with Hoechst dye. Yeast images were collected at 90x magnification on a Nikon Eclipse Te2000-E microscope and processed using ImageJ software. All experiments were repeated at least three times, and representative images are shown.

### Filter Retention Assays

Yeast were grown and induced as for immunoblotting. Following induction, cultures were normalized to A_600nm_=0.5, and 5 mL of cells were collected and washed with sterile water. Cells were resuspended in 500 uL spheroplasting solution (1.2M D-sorbitol, 0.5mM MgCl_2_, 20mM Tris, 50mM β-mercaptoethanol, 0.5 mg/mL Zymolyase 100T, pH 7.5) for 1h at 30ºC with light shaking. Spheroplasts were pelleted by centrifugation at 500x RCF for 5min, and supernatant was discarded. Samples were the resuspended in 100uL lysis buffer (100 mM Tris, pH 7.5, 500 mM NaCl, 5mM MgCl_2_, 10mM β-mercaptoethanol, 0.5% Triton, and 1% yeast Protease Inhibitor cocktail). Samples were vortexed at high speed for 1min and then incubated at room temperature for 10min. Cells were then flash frozen in nitrogen, thawed at room temperature, 33uL of sample buffer (2X TAE, 20% glycerol, 10% β-mercaptoethanol, and 0.0025% bromophenol blue) was added, followed by incubation for 5 minutes at room temperature. 15uL of each extract was applied to a non-binding cellulose acetate membrane using a Minifold I 96-well spot-blot array system (GE). The membrane was then washed with PBST and the remaining sample was diluted in PBST (3 sample:1 PBST) and 5uL was applied onto a nitrocellulose membrane. After both membranes were allowed to dry, they were then re-wetted in PBST for 10min. Membranes were blocked and imaged as described for immunoblotting. Bound protein was quantified using ImageStudio Lite software (LICOR). The reported CA/NC ratio is the ratio of signal on the cellulose acetate (CA) and nitrocellulose (NC) membrane. Values were normalized to a vector control run in parallel.

### Preparation of α-synuclein PFFs

Plasmid for expression of α-synuclein was from Peter Lansbury[31]. α-Synuclein was expressed in *E. coli* BL21-DE3-RIL cells (Invitrogen), where expression was induced at OD_600_=0.6 with 1mM IPTG for 2h at 37ºC. Cell pellets were resuspended in osmotic shock buffer (30mM Tris, pH 7.2, 2mM EDTA, 40% sucrose) and incubated for 10 min at room temperature with vortexing. The lysate was then cleared by centrifugation. Nucleic acids were then removed by streptomycin sulfate precipitation (10mg/mL final concentration) and cleared by centrifugation. The supernatant was then boiled for 10 min and the soluble protein was loaded on to a bed of DEAE sepharose beads for anion-exchange batch purification. The beads were washed with wash buffer (20mM Tris, pH 8.0, 1mM EDTA) and then eluted with elution buffer (20mM Tris, pH 8.0, 300mM NaCl, 1mM EDTA). The eluted protein was then dialyzed against α-Syn fibrillization buffer (20mM Tris-HCl, pH 8.0, 100mM NaCl). Endotoxin was removed using a High-Capacity Endotoxin Removal Spin Column (Pierce) and confirmed using a Charles River Endosafe Cartridge. The protein was then flash frozen and stored at -80ºC until use.

To prepare α-Syn PFFs, monomer was thawed and passed through a 0.2μm syringe filter. The protein was then diluted to 5mg/mL in fibrillization buffer and incubated at 37ºC with agitation at 1,500 rpm in an Eppendorf Thermomixer for 7 days. The resulting mixture was centrifuged at 15,000 rpm for 30 min at room temperature. The supernatant was then removed and fibrils were resuspended to achieve 5mg/mL fibrils, as measured by a BCA assay. Fibrils were then flash frozen and stored at -80ºC until use.

### HEK293T cell culture and FRET

HEK293T biosensor cells (HEK293T-α-syn-CFP/α-syn-YFP) were obtained from David Holtzman[26]. Cells were grown in Dulbecco’s modified high glucose Eagle’s medium (DMEM) supplemented with 10% fetal bovine serum (FBS), and 1% penicillin/streptomycin. For FRET seeding assays, the biosensor cells were plated in 96-well plates at a density of 35K cells per well. 24h following plating, PFFs were applied. Here, α-Syn or Cy5-KFE_8_ fibrils were diluted in fibrillization buffer to the appropriate concentration and sonicated in a cup horn water bath sonicator for 3 min. The fibrils were then mixed with Lipofectamine 3000 (Invitrogen) at 0.5μL Lipofectamine per well, incubated for 10 min at room temperature, and applied to the biosensor cells. After 48 h, cells were trypsinized, transferred to a 96-well plate, and fixed with 4% paraformaldehyde for 15 min at 4ºC in the dark. Cells were then resuspended in 150μL chilled MACSQuant Flow Running buffer for analysis in a MACSQuant VYB flow cytometer. Fluorescence was measured using settings for CFP (ex: 405nm, em: 450, bw: 50nm), YFP (ex: 488nm, em: 525, bw: 50nm), FRET (ex: 405nm, em: 525, bw: 50nm), and Cy5 (ex: 561nm, em: 661, bw: 20nm). Fluorescence compensation was performed with control cell lines each time prior to sample analysis. FRET signal was used to distinguish cells with α-Syn aggregation from cells without α-Syn aggregation. Integrated FRET density was calculated by multiplying the percent cells with FRET signal by median FRET intensity of the FRET-positive cells. To quantify KFE_8_ fibril internalization, a bivariate plot of Cy5 vs CFP was created to introduce a polygon gate to exclude all of the Cy5-negative cells treated with only lipofectamine/buffer and to include the Cy5-positive cells treated with FKE_8_ fibrils. The quantification for KFE_8_ internalization was based on percent Cy5-positive cells. All data analysis was performed with FlowJo V10 software. Images were collected at 20x magnification on a Nikon Eclipse Te2000-E microscope and processed using ImageJ software.

### Generation of Lentiviral Plasmids

293LTV cells were kind gift from Dr. Kian Lim, Washington University in St. Louis. A20 cells were kind gift from Dr. John F. DiPersio, Washington University in St. Louis [32]. pLenti CMV GFP Puro (658-5) was a gift from Eric Campeau & Paul Kaufman (RRID:Addgene_17448)[33]. Packaging pMDLg/pRRE (Addgene plasmid #12251), envelope pMD2.G (Addgene plasmid #12259), and regulatory plasmids pRSV-Rev (Addgene plasmid # 12253) were a gift from Didier Trono[34]. The alpha-synuclein tagged GFP or KFE8_32_ tagged GFP constructs were derived by sub-cloning services from Genescript USA Inc., into pLenti backbone by replacing GFP with αSyn-GFP or KFE_32_-GFP.

### Animals and Immunization

All experimental procedures were approved by the Institutional Animal Care and Use Committee of Washington University in St. Louis. 20μg (100μL of PBS) of pLenti-GFP or αSyn-GFP or KFE_32_-GFP plasmids were injected intramuscularly into 6-8 wk old female BALBC mice three times with 14 days between immunizations. Blood was drawn 10 days after the last dose and sera was collected for ELISA. Spleens were harvested and splenocyte cultures were prepared according to published protocols and stained with H-2Kd restricted GFP_200-208_ tetramer to assess cellular immune responses. Splenocytes (10^6^/well) were also stimulated in vitro with GFP (2 μg/ml) for 5 days and the supernatant collected for analysis of cytokines.

### Antibody titers and isotyping

High binding ELISA plates were coated with 1 μg/mL GFP in PBS and incubated overnight at 4°C. The wells were blocked for 1h at RT with 1% BSA in PBST buffer (PBS with 0.05% Tween20, 100 μL/well). Serial dilutions of the sera were prepared in the blocking buffer and added to the wells (100 μL/well for 1h at RT) followed by secondary HRP-conjugated goat anti-mouse IgG (1:5000, 100 μL/well) for 30 min. For isotyping, Ig isotype antibody IgG1, IgG2b, IgG2c, IgG3, IgM, or IgE, were added (1:1000, 100 μL/well for 1h at RT) after serum dilutions and before the secondary antibody. Plates were washed 3-5 times between each coating step. The assay was developed using TMB solution (100 μL/well) for 15 min and the reaction was quenched with 1 M phosphoric acid (100 μL/well). Absorbance was recorded at 450 nm using a Biotek Synergy microplate reader.

### Cellular Immune Responses

Induction of GFP-specific T cell response was quantified by measuring the tetramer positive CD8^+^T cells in spleen of vaccinated mice via flow cytometry. Briefly, splenocytes from vaccinated mice were prepared following RBC lysis of whole spleen suspensions as reported previously. Cells were stained with Zombie green fixable viability dye for 15 minutes, washed in once with FACS buffer, and stained with H-2Kd restricted GFP_200-208_ (HYLSTQSAL) tetramer (PE, # TS-M525-I, MBL Inc.) followed by CD3 (PE-Cy7) and CD8 (APC) staining. Total 10^6^ events were recorded on NovoCyte instrument (ACEA Biosciences, USA.) and the data were analyzed by FlowJo v.10 software (FlowJo, LLC). To assess cytokine production, splenocytes (10^6^/well) were plated in 24-well plates and treated with GFP (2 μg/ml for 5 days or with PBS as control). Supernatant was collected and cytokines were quantified using mouse cytokine & chemokine convenience 6-Plex kit (IL-2, IL-4, IL-5, IL-10, IFN-γ and TNF-α; # LXSAMSM-06, R&D systems) using Luminex xMAP according to manufacturers protocols.

## RESULTS

### Testing the toxicity and clearance mechanism of peptide nanofiber constructs in yeast

*Saccharomyces cerevisiae*, Baker’s yeast, has long been a useful model system for studying biology. Though simple, yeast is highly genetically tractable, and genetic pathways are highly conserved between yeast and humans[35]. Yeast has been shown to be an excellent model system for studying complex human diseases including ALS/FTD, Parkinson’s disease, Alzheimer’s disease, diabetes, and sarcoma[36-43]. We therefore sought to study the properties of the peptide nanofiber constructs in yeast, compare their properties to those of disease-associated amyloid and amyloid-like proteins, and follow their mechanism of clearance. To test the effects of intracellular expression of the nanofiber constructs, we generated plasmids encoding 1, 2, or 4 repeats of the sequence FKFEFKFE for expression in yeast. Each repeat was separated by a flexible glycine-serine linker, and to monitor expression and localization, we fused each construct to eGFP. We used the 423GAL plasmid to drive galactose-inducible expression of the genes, and we expressed KFE_8_, KFE_16_, and KFE_32_, along with a GFP control. We also included the amyloid-like protein TDP-43, which has been implicated in ALS/FTD and has been shown to be highly toxic and aggregate in yeast[44]. As expected, TDP-43 was highly toxic in yeast, while KFE_8_ and KFE_16_ were nontoxic. KFE_32_ was just slightly toxic (**Fig 1A**). Expression of each of the constructs was confirmed via immunoblotting, and we also observed the expected increase in molecular weight corresponding to increased length of repeats (**Fig 1B**).

**Figure 1.**
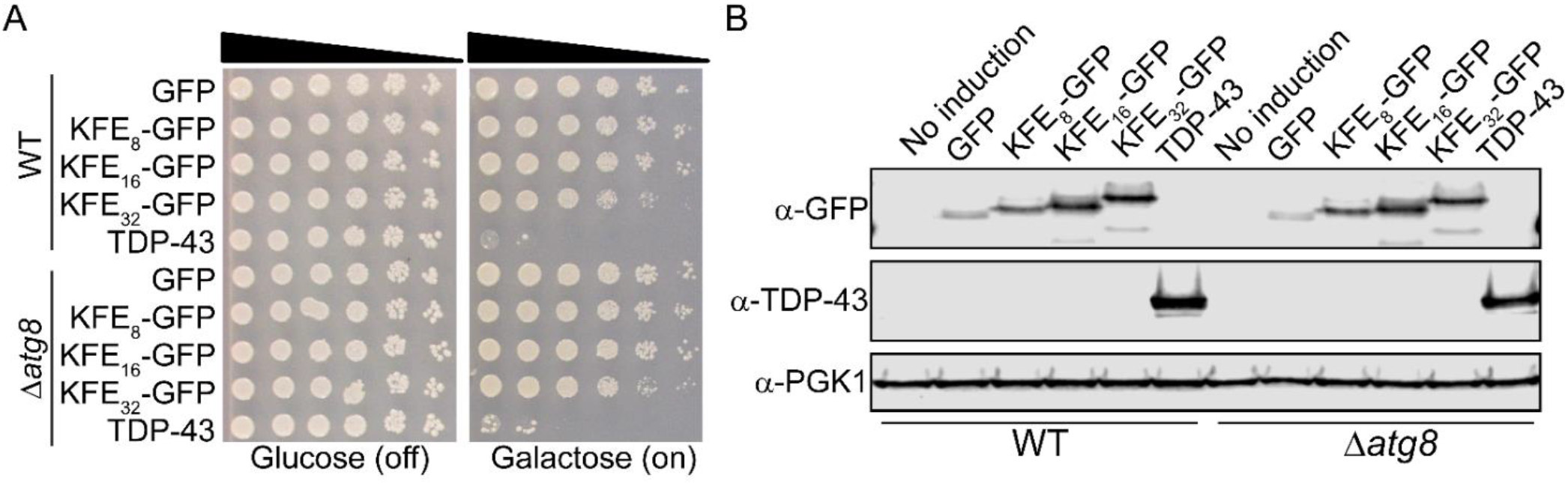
KFE_8_-GFP fusions are nontoxic in yeast. (A) BY 4741 and BY Δ*atg8* yeast was transformed with galactose-inducible KFE_8_-GFP fusions of varying lengths along with a GFP and TDP-43 control. The strains were serially diluted 5-fold and spotted onto glucose (off) or galactose (on) media. (B) Strains from (A) were induced for 5 h, lysed, and immunoblotted. PGK1 serves as a loading control.

It has been demonstrated that these peptide nanofibers engage the autophagy machinery in mammalian cells[45]. Such off-target effects could prove problematic for ultimate therapeutic applications of the nanofibers, particularly in elderly populations known to have impaired autophagy, though toxicity has not been noted in mammalian cell studies. Therefore to validate our yeast model, we sought to assess if these proteins also engage autophagy in yeast. We assessed the toxicity of each of the fusions in Δ*atg8* yeast and found that toxicity was not modified in this background as compared to WT yeast (**Fig 1A**). We therefore conclude that the peptide nanofibers are not toxic and toxicity is not exacerbated under autophagy-deficient conditions.

### Peptide nanofibers form insoluble inclusions in yeast

Amyloid proteins are known to form insoluble inclusions that are resistant to boiling and other denaturing conditions. To determine if the KFE repeats formed insoluble species in yeast, we employed a filter retention assay[46]. Here, following induction of expression as for immunoblotting, extracts are applied to a non-binding cellulose acetate and a binding nitrocellulose membrane in parallel. Larger insoluble species will be retained on the cellulose acetate membrane while smaller species will pass through the membrane. Proteins that form highly stable amyloid conformers will be resistant to high concentrations of denaturant and boiling. Here, we found that each of the KFE_8_ fusions strongly bound to the cellulose acetate membrane as compared to the GFP control (**Fig 2A**). Following boiling, all samples passed through the membrane. The ratio of material retained on the two membranes was quantified and we observed a predicted increase in retention on the cellulose acetate membrane that correlates with increased length of the construct (**Fig 2B**). We therefore conclude that the peptide nanofiber constructs assemble into stable, insoluble species, but that these species are not as stable as amyloid species and can be disrupted at high temperatures.

**Figure 2.**
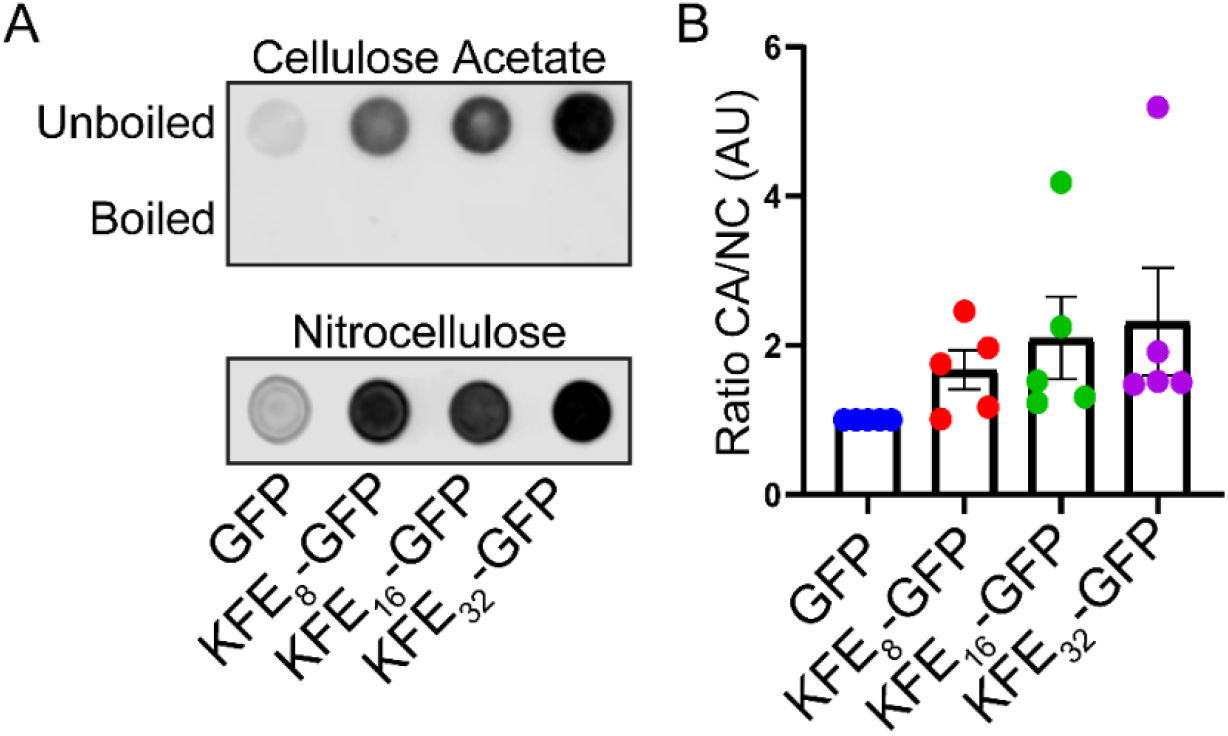
KFE_8_-GFP fusions are insoluble in yeast. (A) Filter retention assays of the three KFE_8_-GFP fusions expressed in WT yeast. Cells were induced for 5h, lysed, and passed in duplicate through a nonbinding cellulose acetate membrane and a binding nitrocellulose control membrane. Samples were also boiled and passed through the cellulose acetate membrane. Peptides were then detected by immunoblotting for the GFP epitope. Experiments were performed in triplicate with independent transformations, representative results are shown. (B) The ratio of binding to the cellulose acetate (CA) to nitrocellulose (NC) membrane for the unboiled samples was quantified and normalized to the vector control. Dots show biological replicates, bars show means, and error bars show SEM (N=5). A one way ANOVA with Tukey’s multiple comparison test was used to assess differences among each of the strains and no significance was found.

### Peptide nanofibers form foci in yeast that are cleared by autophagy

We next sought to visualize the properties of the peptide nanofibers in yeast using fluorescence microscopy. We studied each of the strains and found that expression of each of the six strains elicited formation of foci in yeast (**Fig 3A**). We quantified these effects and found no significant differences in the number of cells harboring foci that correlated with length of the KFE_8_ repeat, although the foci did appear larger in strains expressing KFE_16_ and KFE_32_ as compared to KFE_8_. We also observed that many cells in the WT background displayed GFP fluorescence in the vacuole (**Fig 3B**), while vacuolar accumulation was not observed in the Δ*atg8* background. We quantified these effects and saw a subtle, but consistent, increase in the number of cells with foci in the Δ*atg8* background as compared to the WT background (**Fig 3C**). Based on this increased number of foci in the autophagy-deficient strain, and the accumulation of vacuolar KFE, we conclude that the nanofibers are trafficked through the autophagy pathway in yeast, as they are in mammalian cells, yet this does not trigger toxicity.

**Figure 3.**
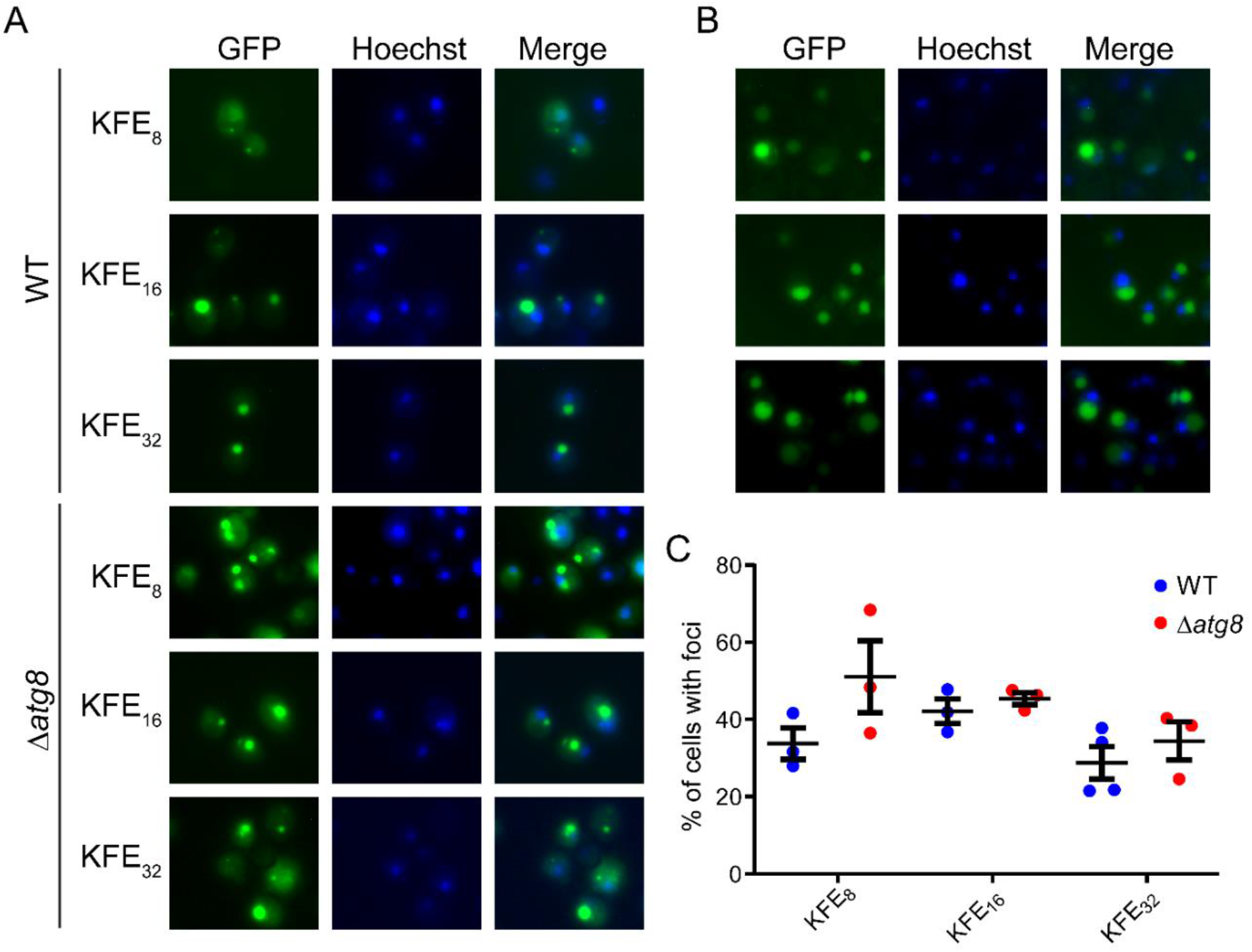
KFE_8_-GFP fusions form foci in yeast. (A) Fluorescence microscopy of the three KFE_8_-GFP fusions expressed in WT and Δ*atg8* yeast. Strains were induced for 15 h, stained with Hoechst dye to visualize nuclei (blue), and imaged. Foci formed in each strain, with no clear differences among the strains. Representative images are shown. (B) Additional images of strains shown in (A). In some cells in the WT background, vacuolar accumulation of GFP was observed. No vacuolar accumulation of GFP was observed for cells in the Δ*atg8* background. (C) Quantification of microscopy experiments shown in (A). Error bars show SEM. A one way ANOVA with Tukey’s multiple comparison test was used to assess differences among each of the strains and no significant differences were found.

### Peptide nanofibers cannot cross-seed aggregation of α-synuclein

Amyloid fibrils are known to spontaneously enter cells and seed aggregation of other copies of monomeric protein. These features underpin the cell-to-cell transmissibility and infectious nature of amyloid, allowing for a small quantity of seed to rapidly accelerate conversion of monomer to adopt the amyloid fibril conformation[47]. Further, cross-seeding is a phenomenon by which amyloid fibrils comprised of one protein can nucleate amyloid formation by a protein of a different sequence. Given the resemblance of the secondary structure of KFE nanofibers to disease-associated amyloid and amyloid-like proteins, it is of crucial importance that the KFE nanofibers be incapable of seeding aggregation of disease-associated amyloid proteins. To test this idea, we employed HEK293T α-synuclein biosensor cells[25, 26]. Here, one copy of α-synuclein is fused to cyan fluorescent protein (CFP) while another copy is fused to yellow fluorescent protein (YFP). CFP and YFP function as a pair of fluorophores that can readily undergo fluorescence resonance energy transfer (FRET) when in close proximity, such as upon aggregation. When pre-formed α-synuclein fibrils (PFFs) are added to cell culture medium, the PFFs are internalized by the HEK cells and trigger aggregation of α-synuclein which is quantified by measuring the FRET signal by flow cytometry (**Fig 4A**)[26].

**Figure 4.**
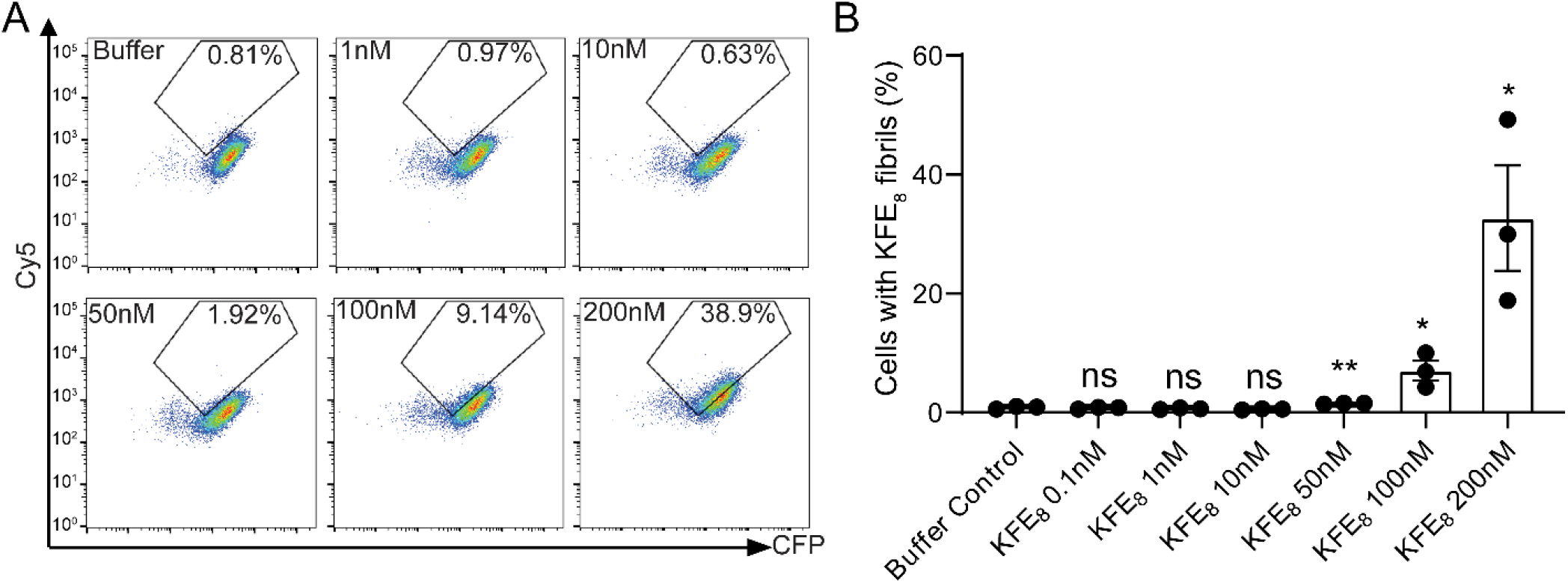
KFE_8_ nanofibers are internalized by HEK biosensor cells. (A) Cy5-KFE_8_ nanofibers were applied to HEK biosensor cells and internalization was monitored by flow cytometry. (B) Quantification of cells from A based on Cy5 signal. Values were compared to buffer control using a series of two-tailed t-tests (N=3, biological replicates are shown as dots, bars represent means ± SEM, *p<0.05, **p<0.01).

To test if the KFE nanofibers can seed α-synuclein biosensor cells, we first confirmed uptake of the fibrils by flow cytometry using Cy5 labeled KFE fibrils. Internalization of the nanofibers was robust, with approximately 10% of cells displaying a Cy5 positive signal on addition of 100nM fibrils. When the concentration was increased to 200nM, nearly 40% of cells were Cy5 positive (**Fig 4**). We next assessed seeding. Here, application of just 50nM α-syn PFFs is sufficient to trigger a robust FRET signal (**Fig 5**), with over 25% of cells displaying FRET. When Cy5-KFE_8_ preformed fibrils are applied instead of α-syn PFFs we observe no increase in FRET (**Fig 5**). We increased the concentration of Cy5-KFE_8_ to as high as 200nM and still observed no FRET signal (**Fig 5**), confirming that the nanofibers do not seed α-synuclein aggregation. As a final test, we applied 10μM KFE_8_ nanofibers without the Cy5 tag to ensure there was no interference with the FRET pair and still observed no seeding (**Fig 5**). To further corroborate these results, we imaged these cells using fluorescence microscopy. Again we observe an abundance of foci upon application of αSyn PFFs, while no foci are observed upon application of the KFE_8_ nanofibers (**Fig 6**). Thus we conclude that the KFE nanofibers can enter cells similarly to amyloid fibrils, but they cannot cross-seed α-syn.

**Figure 5.**
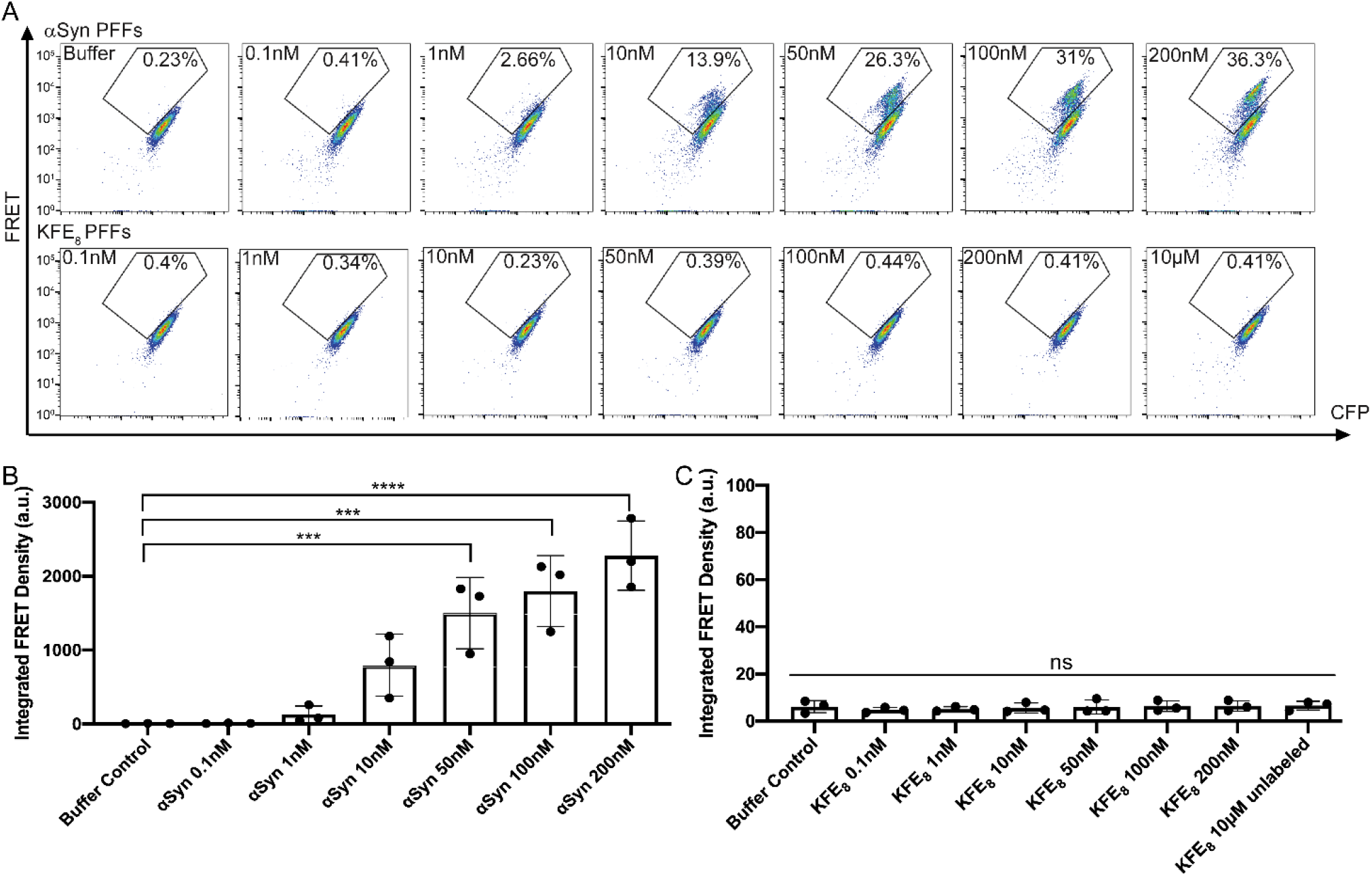
KFE_8_ nanofibers do not cross-seed α-synuclein. (A) HEK α-synuclein biosensor cells were seeded with αSyn PFFs (top row) or KFE_8_ PFFs (bottom row) at the indicated PFF concentrations. Seeding of αSyn was monitored by flow cytometry. (B) Integrated FRET density for the αSyn PFFs was quantified from the experiments in A. (C) Integrated FRET density for the KFE_8_ PFFs were quantified from the experiments in A. Values in B and C are compared to buffer treatment using a one-way ANOVA with a Dunnett’s multiple comparisons test (N = 3, biological replicates are shown as dots, bars represent means ± SEM, ***p < 0.001, ****p<0.0001).

**Figure 6.**
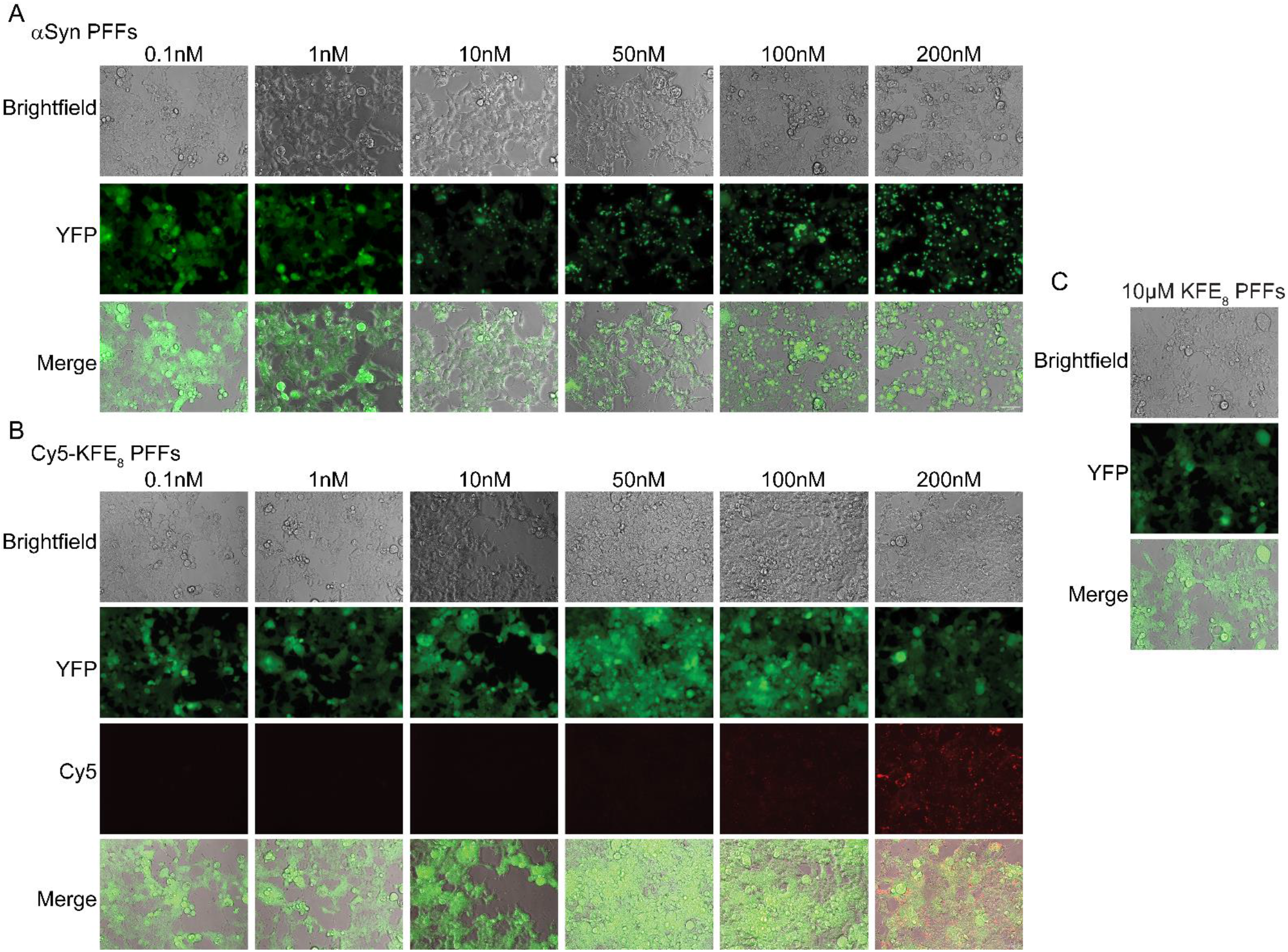
KFE_8_ nanofibers do not cross-seed α-synuclein. (A) α-Syn PFFs were added to biosensor cells at the indicated concentration and imaged. (B) Experiments were performed as in A but with Cy5-KFE_8_ PFFs instead of α-Syn. (C) Experiments were performed as in B but with unlabeled KFE_8_ at higher concentration. Scale bar = 20μm, scaling is the same in all images.

### Vaccination with plasmids encoding KFE_32_-GFP elicits a robust antibody response

DNA-based vaccines have many advantageous features as compared to synthetic peptide vaccine strategies such as decreased cost, improved stability, and the ability to construct longer sequences than can be accomplished using solid-phase peptide synthesis. Yet, synthesis in the host may also lead to different assembly patterns and architectures than when synthesized in vitro. Inspired by the striking similarities we have observed with synthetic KFE_8_ fibrils and upon expression in yeast, we next explored the capacity for these materials to be encoded in a DNA-based vaccine. Here, we primed and boosted groups of Balb/c mice with plasmids encoding GFP, α-Syn-GFP or KFE_32_-GFP and assayed the levels of anti-GFP antibodies in the sera. Data indicated that mice vaccinated with the KFE_32_-GFP construct produced significantly higher levels of anti-GFP antibodies as compared to soluble GFP or GFP-α-Syn conjugates (**Fig 7A**). This difference can be ascribed to the monomeric or oligomeric nature of soluble GFP or α-Syn-GFP, respectively, whereas KFE_32_-GFP could potentially fibrillize to form higher molecular weight, high aspect ratio nanofibrils that mimic danger signals and activate the immune system. No significant differences were detected between mice vaccinated with control GFP or α-Syn-GFP plasmids at all serum dilutions tested. In order to determine the nature of the immune response, the isotypes of the responding antibodies were evaluated. For all constructs IgG1, IgG2b, and IgG2c, IgG3, and IgM were produced in similar quantities suggesting a balanced response with strong bias towards a Th1-or Th2-type phenotype (**Fig. 7B**). Interestingly, levels of IgE were significantly higher in mice vaccinated with KFE_32_-GFP plasmid compared to controls (**Fig 7B**). These findings are exciting as they demonstrate that nanofiber vaccines can be produced upon expression driven by a plasmid in a host, allowing for more complex constructions than is possible with materials produced by in vitro synthesis. Further, prior studies have demonstrated that vaccination with synthetic peptide nanofibers generally induces a Th2-bias with dominant IgG1 production without any detectable allergic IgE. These data demonstrate that DNA vaccination with model antigens conjugated to KFE_8_ repeats induces robust antibody responses similar to peptide-based vaccines but with distinct Th1/Th2 profiles.

**Figure 7.**
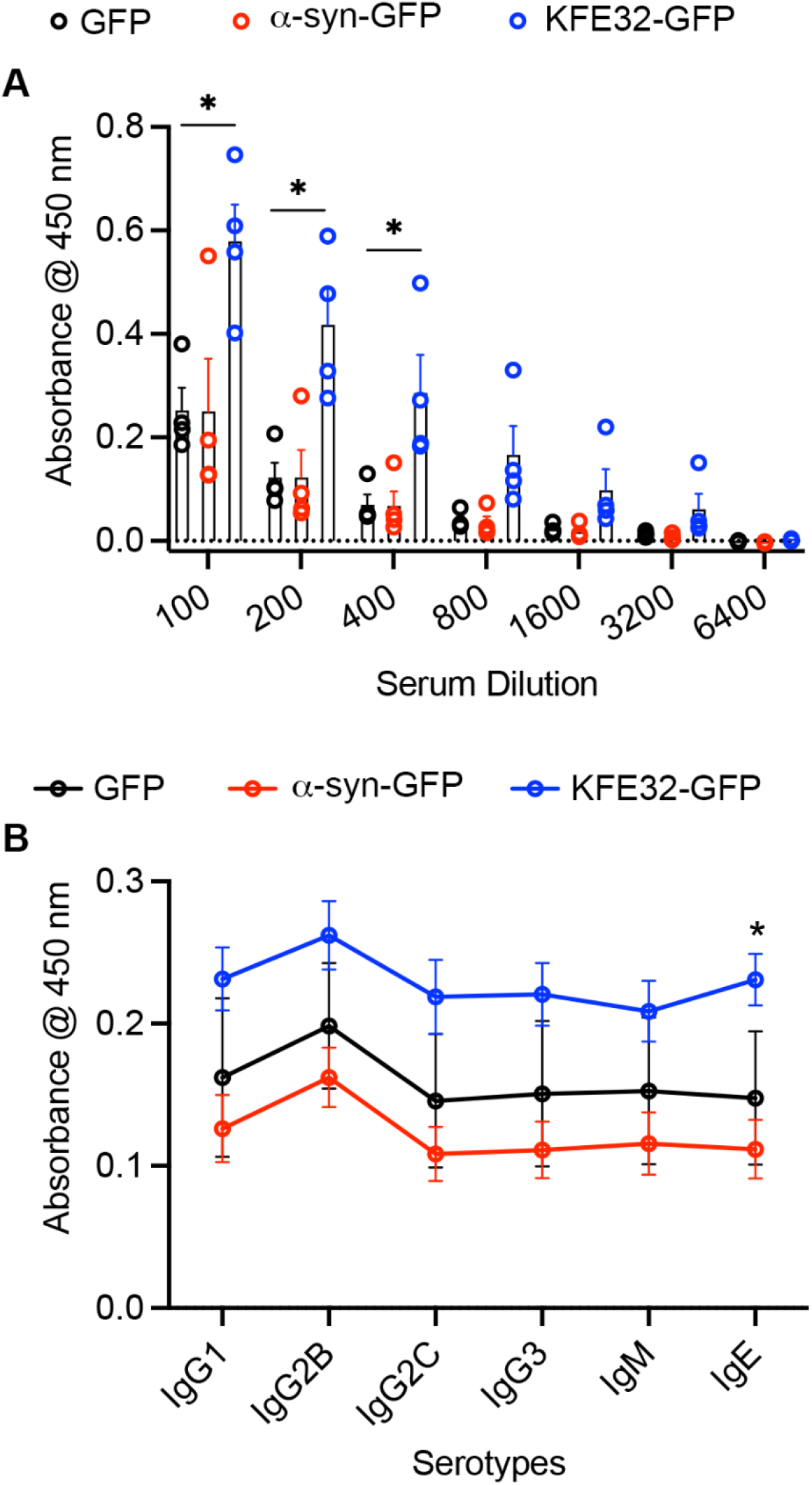
DNA vaccination with KFE_32_-GFP constructs elicits robust antibody responses. **(A)** Prime-boost with KFE_32_-GFP construct strongly adjuvanted antibody responses against GFP and at significantly higher levels compared to αsyn-GFP or soluble GFP. Similar titers of total IgG were raised against a-syn-GFP fusion or soluble GFP. (B) Antibody isotypes IgG1, IgG2b, IgG2c, IgG3, IgM, IgE in sera from immunized mice. IgE was significantly higher in the sera of mice vaccinated with KFE_32_-GFP. *p < 0.01 by ANOVA with Tukey HSD post hoc testing, compared between groups as indicated.

### Vaccination with plasmids encoding KFE32-GFP elicits a robust T cell response

Immunization with DNA is known to be an effective inducer of cellular immunity. Having demonstrated that our constructs elicit a robust antibody response (**Fig 7**), we next sought to evaluate the ability of GFP, α-Syn-GFP or KFE_32_-GFP constructs to potentiate the production of antigen-specific CD8^+^T cells. Spleens from vaccinated mice were used in antigen recall assays and the frequency of tetramer+ CD8^+^T cells was quantified using flow cytometry. Surprisingly, higher levels of antigen-specific cells were detected in mice receiving the soluble GFP protein construct compared to α-Syn or KFE_8_ fusions, though this did not reach statistical significance (**Fig 8**). Importantly, both α-Syn and KFE_8_ fusion constructs were found to elicit equivalent levels of antigen-specific CD8^+^T cells (**Fig 8**). The augmentation of antibody responses, isotype class-switching, and tetramer+ CD8^+^T cells also point to the possibility that KFE_32_-GFP could efficiently trigger CD4^+^T cell mediated helper immunity. To investigate the involvement of T cell help, splenocytes from immunized mice were challenged in vitro with soluble GFP (cognate antigen), and the production of interferon-γ (IFN-γ), tumor necrosis factor-α (TNF-α) interleukin-2 (IL-2), interleukin-4 (IL-4), interleukin-5 (IL-5), and interleukin-10 (IL-10) was measured using a multiplex assay. These cytokines were selected to provide measures of either a Th1 response (IFN-γ, TNF-α, and IL-2) or a Th2 response (IL-4, IL-5, IL-10). Data indicated significantly higher production of all cytokines in splenocyte cultures of mice vaccinated with soluble GFP, α-Syn-GFP, or KFE_32_-GFP constructs compared to controls (splenocytes from the same mice that did not receive the cognate GFP stimulus) (**Fig 9**). This finding also supports the antibody isotyping data where diverse serotypes were detected without a dominant Th1/Th2 bias.

**Figure 8.**
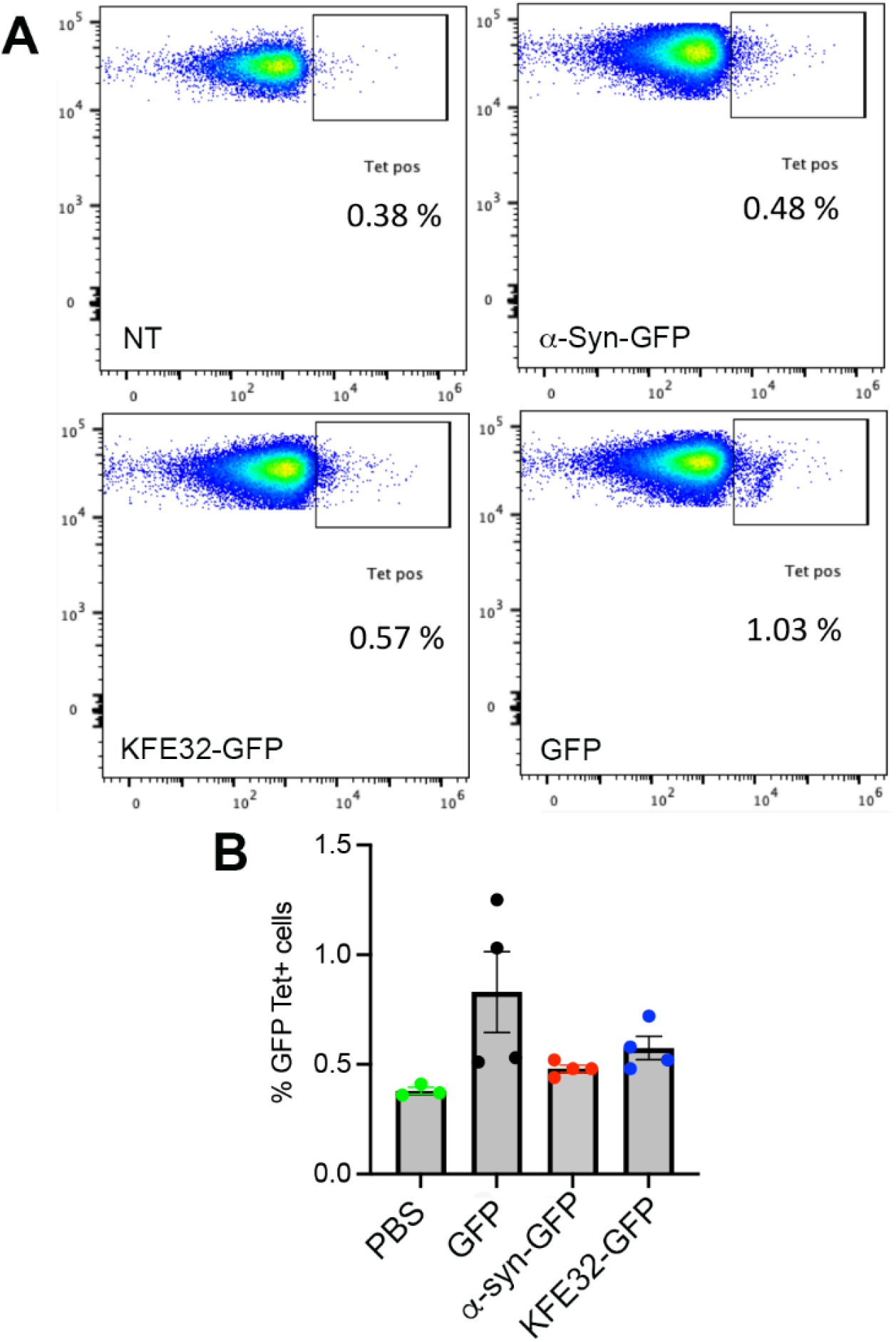
CD8^+^T cell responses in mice vaccinated with KFE_32_-GFP or controls. (A) Schematic showing immunization regime and timeline. (B) Representative flow cytometry plots showing production of tetramer^+^ CD8^+^ T cells in the spleens of mice immunized with soluble GFP, αsyn-GFP, or KFE_32_-GFP. (C) Cumulative bar graph from (B) showing the percentage of tetramer^+^ CD8^+^ T cells (*n* = 4 mice per group).

**Figure 9.**
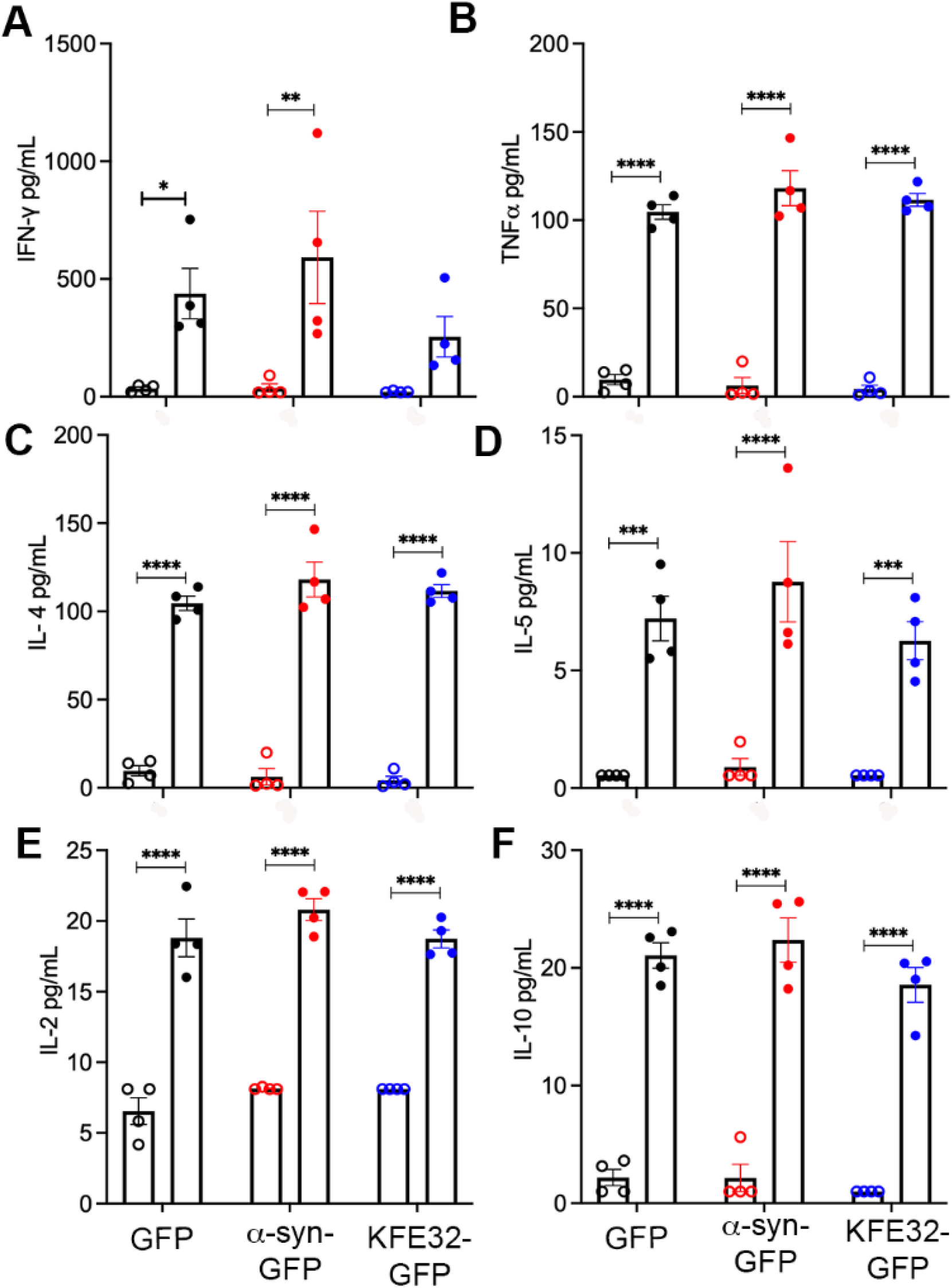
DNA vaccination with KFE_32_-GFP elicits a balanced Th1/Th2 response. (A) Production of Th1 cytokines (A) IFN-γ, (B) TNF-α (E) IL-2 and Th2 cytokines (C) IL-4 (D) IL-5 and (F) IL-10 in unstimulated (open circle) or stimulated (solid circle) splenocyte cultures from mice vaccinated with soluble GFP, αSyn-GFP, or KFE_32_-GFP constructs. *p < 0.05, **p < 0.01, ***p< 0.001, ****p <0.0001 compared with corresponding unchallenged control, by ANOVA with Sidak’s multiple comparisons test.

## DISCUSSION

A crucial challenge in vaccine development is to achieve a strong immunogenic response while also ensuring safety[48]. Most currently used vaccines rely on co-administration with adjuvants to enhance, maintain, and direct the adaptive immune response. In the U.S., only aluminum-based adjuvants are currently licensed for human use, though a new generation of adjuvants is under development[49]. These adjuvants are chemically heterogeneous mixtures of plant- or pathogen-derived products, formulations of mineral salts, or emulsions, all of which which suffer from poor chemical definition and are associated with toxicity. These limitations have motivated the pursuit of adjuvant-free vaccination technologies.

To enhance vaccine-based immunotherapies to combat infections, cancer, and other conditions, biomaterials are being heavily explored with the goal of improving vaccine safety and efficacy[50]. A prominent strategy, inspired by viruses, is to leverage macromolecular self-assembly which facilitates the multivalent presentation of antigens, significantly enhancing immunogenicity. Recently, peptides that self-assemble into specific nanoarchitectures have been pursued as modular and chemically defined platforms for vaccine development[4]. In particular, ‘amyloid-inspired’ peptide nanofibers that adopt a cross-β structure have been demonstrated to be effective for efficient delivery of vaccines and immunotherapies in multiple preclinical models of infectious and non-infectious diseases[4]. However, several unresolved challenges still remain and must be addressed to accelerate translational efforts. A key hurdle is the fact that the safety of using cross-β sheet rich amyloid-like structures must be carefully studied[4].

The field of peptide self-assembly was motivated by initial attempts to gain molecular and structural information on misfolded states of proteins involved in neurodegenerative diseases. Identification of minimal peptide fragments capable of emulating the behavior of the parent protein was used as a reductionist approach to glean structural information on these systems and understand disease pathogenesis. Unexpectedly, these studies also suggested that very small peptides are capable of self-assembly. Minimal sequences derived from natural proteins such as islet amyloid polypeptide (FGAILK), human calcitonin (DNFKFA), and Aβ_1-42_ (KLVFFA) have been reported[51-53]. Reches and Gazit first identified the diphenylalanine (FF) peptide as the shortest motif able to self-assemble from the full length amyloid beta peptide (Aβ_1-42_)[54]. A second family of cross-β rich peptides comprised of amphipathic sequences with strictly alternating apolar and charged residues, such as the protopeptide EAK [Ac-(AEAEAKAK)2-NH2], were first identified in *S. cerevisiae*[55]. Because these peptides all self assemble into structures resembling amyloid fibrils, and amyloid underpins the pathogenesis of several neurodegenerative diseases, understanding the amyloidogenic potential of cross-β rich peptide biomaterials intended for vaccine development or other *in vivo* applications is paramount for their continued success. Computational and spectroscopic efforts in structural biology have identified that cross-β rich peptide aggregates are characterized by remarkable molecular and structural complexities that correlate with toxicity. Importantly, the outcomes of these studies have been exploited to design peptide inhibitors of amyloid-aggregation and cross-β rich peptide fibrils have also been reported to be capable of triggering innate immune pathways for the design of vaccines [8, 56-58]. Also, emerging insights into the functional roles of amyloid and amyloid-like proteins have challenged the notion that all amyloid species are inherently toxic, and crucial differences that distinguish pathological, inconsequential, and functional aggregation are not yet fully understood.

In this study we utilized the self-assembling peptide KFE_8_ (FKFEFKFE). This peptide sequence is not only Phe rich but also amphipathic with alternating apolar and charged residues. Association of the two apolar faces generates the basic cross-β motif that constitutes the backbone of the fibrils. The structure, molecular packing, and assembly of KFE_8_ have been extensively characterized using spectroscopic and computational studies[7, 16, 17]. Further, we have demonstrated the ability of KFE_8_ to act as an immune adjuvant in multiple preclinical models of disease[4]. Therefore, we anticipate that, barring safety concerns, the KFE_8_ sequence may have broad translational utility. Here, to further the translational potential of amyloid-inspired peptide nanofibers, we have investigated the properties of the self-assembling peptide KFE_8_. First, we leveraged the genetic tractability of *S. cerevisiae*, Baker’s yeast, to characterize varying lengths of this repeat. We find that KFE_8_ repeats have distinct properties from those of amyloid and amyloid-like proteins. While KFE_8_ forms characteristic insoluble inclusions, expression of these materials is non-toxic. We have previously reported that KFE_8_ fibrils can enter cells via endocytic mechanisms, and that the adjuvanting potential of peptide nanofibers is due to their ability to engage key components of the autophagy pathway[45]. Triggering autophagy is important, because this significantly improves responses to influenza vaccines in the elderly and the BCG vaccine in infants[59]. However, amyloid species are also cleared via autophagy, and evidence suggests that impairment of autophagy in the elderly may be linked to increased susceptibility to neurodegeneration with aging[60]. Here, we find that just as these peptides are cleared via autophagy in mammalian cells, they are also cleared via autophagy in yeast. Promisingly, even in an autophagy-impaired strain, expression of the KFE_8_ sequences remains non-toxic in yeast. The ability to study these biomaterials upon expression in yeast will be a useful platform in further characterization of these materials. For instance, this yeast expression platform can be used to conduct a genome-wide screen to investigate if the nanofibers engage or interact with other proteins in the cell, and to investigate and/or identify possible off-target effects.

Accumulating evidence implicates transcellular propagation of amyloid species, or seeds, as a mechanism for disease progression in neurodegeneration. Amyloid species are also known to be capable of ‘cross-seeding’, whereby amyloid fibrils comprised of one protein can initiate the amyloid cascade of another amyloidogenic protein. This is thought to be due to the highly conserved structural features of amyloid and pre-amyloid, which are highlighted by the findings that a single antibody can cross-react with pre-amyloid species comprised of a diverse range of monomeric proteins[12]. As such, it is crucial to investigate the capacity of any amyloid-like therapeutics to seed amyloidogenesis of endogenous proteins. Here, we find that while α-syn preformed fibrils can enter the cell and trigger robust seeding of endogenous α-syn, KFE_8_ fibrils can also enter the cell, but they do not trigger seeding of α-syn amyloidogenesis. In future work it will be important to confirm that these fibrils do not cross-seed a broader range of amyloidogenic proteins.

Finally, inspired by our successful expression of the KFE-GFP fusions in yeast, we also explored the development of DNA-based vaccines using the KFE platform. DNA-based vaccines remain an infrequently employed mode of delivery, yet this strategy has great promise. By genetically encoding synthesis of a vaccine, longer sequences can be made than is practical by synthetic routes. Further, DNA is less expensive to generate, easier to modify, and can be stored at ambient temperatures. To investigate these applications, we immunized mice with plasmids encoding KFE_32_-GFP, where GFP serves as a model antigen. Immunization elicited a robust immune response, and the KFE_32_ scaffold induced production of significantly higher levels of anti-GFP antibodies as compared to immunization with plasmid encoding GFP alone. Further, the cytokine profiles from antigen recall assays indicate a balanced Th1/Th2 response. The self-adjuvanting potential observed in the synthetic peptide form of the nanofibers was not diminished when delivered for expression from a DNA plasmid.

In summary, we demonstrate that expression of KFE_8_ results in the formation of insoluble inclusions in yeast similar to those of amyloid and amyloid-like proteins. However, KFE_8_ is not toxic and toxicity is not exacerbated in autophagy-deficient yeast. Further, synthetic KFE_8_ nanofibers do not cross-seed the amyloidogenesis of α-synuclein, even at 1000× higher concentrations compared to α-syn preformed fibrils. In mice, vaccination with plasmids encoding KFE_32_-GFP elicited a robust cellular immune response with a balanced Th1/Th2 cytokine profile, along with generation of antigen-specific CD8^+^T cells. These findings are significant and exciting because, since their serendipitous discovery, several applications based on self-assembling peptide nanofibers have been brought to the market and new sequences are being designed to generate chemically defined and hierarchically organized nanoscale materials. However, to date no studies have investigated the amyloidogenic potential of self-assembling peptides or their utility as DNA vaccines. Our results demonstrate that despite their molecular, structural, and morphological similarity to pathological amyloids, self-assembling peptides have distinct properties, and are promising materials for biomedical applications.

## DATA AVAILABILITY

Data will be made available on request.

## ACKNOWLEDGEMENTS

Our studies were supported by NIH grant R21AG068733 (to J.S.R. and M.E.J.) and R01 AI130278 to J.S.R.

